# Optical trapping of nucleolus reveals viscoelastic properties of nucleoplasm inside mouse germinal vesicle oocytes

**DOI:** 10.1101/2020.03.19.999342

**Authors:** Maria S. Syrchina, Aleksander M. Shakhov, Arseny V. Aybush, Victor A. Nadtochenko

## Abstract

We propose a technique of controlled manipulation with mammalian intracellular bodies by means of optical trapping in order to reveal viscoelastic properties of cell interior. Near infrared laser in the spectral range of tissue transparency was applied to study dynamics of the nucleolus-chromatin complex inside the thermodynamically non-equilibrium system of a mouse oocyte. A nucleolus of germinal vesicle (GV) oocyte as spherical probe was displaced from the equilibrium and its relaxation dynamics was observed. We developed software for subdiffraction tracking of a nucleolus position with lateral resolution up to 3 nm and applied it for different GV-oocyte chromatin configurations. We showed differences in viscoelastic properties within nucleoplasm of NSN-oocytes, visualized by Hoechst 33342 staining. Also, we demonstrate that in germ cells basic biophysical properties of nucleoplasm can be obtained by using optical trapping without disruption and modification of cellular interior.

---

Macromolecules confined in the nuclear envelope adjust their properties in respond to extra- and intracellular signals, providing a resistance of the cell to continuously changing environment. Cell nucleus represents complex organelle composed of polymer bodies of different size and shape, constantly diffusing through the nucleoplasm. Understanding of the mechanical alterations experienced by nucleic acids and proteins within the nucleoplasm is crucial for clarifying genome functioning.

Two general concepts of large-scale nuclear organization were earlier proposed and are still being discussed. First one describes nucleus as a structure with highly compartmentalized inner space supported by nuclear matrix.^1-3^ But there is a large body of evidence suggesting, that nucleus has dynamic, liquid-like, disorganized, viscous environment and includes constantly diffusing and assembling/disassembling nuclear bodies. ^4-7^

Rheology of cellular soft materials is defined by key physical parameters – viscosity, elasticity and creep compliance. These properties of cellular interior can be obtained by particle-tracking measurements. Strategies for estimating intracellular rheology subdivided into active (microparticles are subjected to external forces)^8^ and passive techniques (tracking the Brownian motion of embedded particles or cellular bodies).^9-11^ Methods of particle-tracking microrheology are usually combined with introduction of fluorescent probes (FCS, FRAP);^12-14^ or microbeads^15^ with magnetic^16^ and optical trapping;^17-18^ and cantilever of atomic force microscope,^19^ what makes them indispensable tools for extracting essential biophysical parameters of cellular interior.

Optical trapping gives precise control and positioning of the micron size objects by means of tightly focused and localized laser radiation. Organelles can be employed as microprobes in non-invasive investigation of cellular interior by means of optical trapping.

The global role of oocyte is to accurately transfer genetic material and initiate embryo development. Well known, that mechanical alterations of cell nucleus can provoke deep physiological changes of the whole organism.^20^ Research of nuclear micromechanics is traditionally performed on somatic cells in the context of mitotic cycle, thereby biophysics of intranuclear space during oogenesis at different developmental stages is poorly understood. Complexity of largescale spatial chromatin organization during meiosis is not quite accurately represented by models, traditionally applied for interpretation results of interphase chromatin dynamics (even characterizing the loop-organized chromatin).^21-22^

Mouse germinal vesicle (GV) oocytes are casually divided into two main groups in accordance with chromatin profile inside the nucleus. Chromatin in non-surrounded nucleolus (NSN) oocyte is diffusely distributed in the nucleoplasm, whereas heterochromatin surrounds the nucleolus with a ring in surrounded nucleolus (SN) oocyte.^23^ These states differ at biochemical and morphological levels and, as a result, in the potential for successful resumption of meiosis, fertilization and development.^24^ As an object, we chose mouse oocyte on a specific stage (GV) of meiotic maturation – diplotene, that is not typical for nuclear micromechanics investigation, but is the matter of interest in the field of developmental biology.

To show the differences between micromechanics within the population of NSN-oocytes, tracking of nucleolus motion, induced by the optical trap, was performed. Optical tweezers supplied system with an external force and nucleoli of mouse oocytes were employed as microspheres. Using of nucleolus, as a largest nuclear body of mouse oocyte, excluded pulling of smaller nuclear bodies inside the optical trap. After series of subsequent displacements of nucleolus in several directions, nucleolus-chromatin complex was allowed to relax. Videos were processed with sub-diffraction resolution by custom-made software and characteristic relaxation times were extracted. Chromatin was visualized by Hoechst 33342 dye in order to compare large-scale profile for different oocyte populations.

## RESULTS

Initially, we divided oocytes according to the classification based on chromatin configuration after Hoechst 33342 staining.^25^ Among them, NSN-oocytes (compared with SN-oocytes) displayed a relatively homogenous nucleoplasm, higher range of directions for nucleolus displacements (Fig. 1).

**Figure 1.**
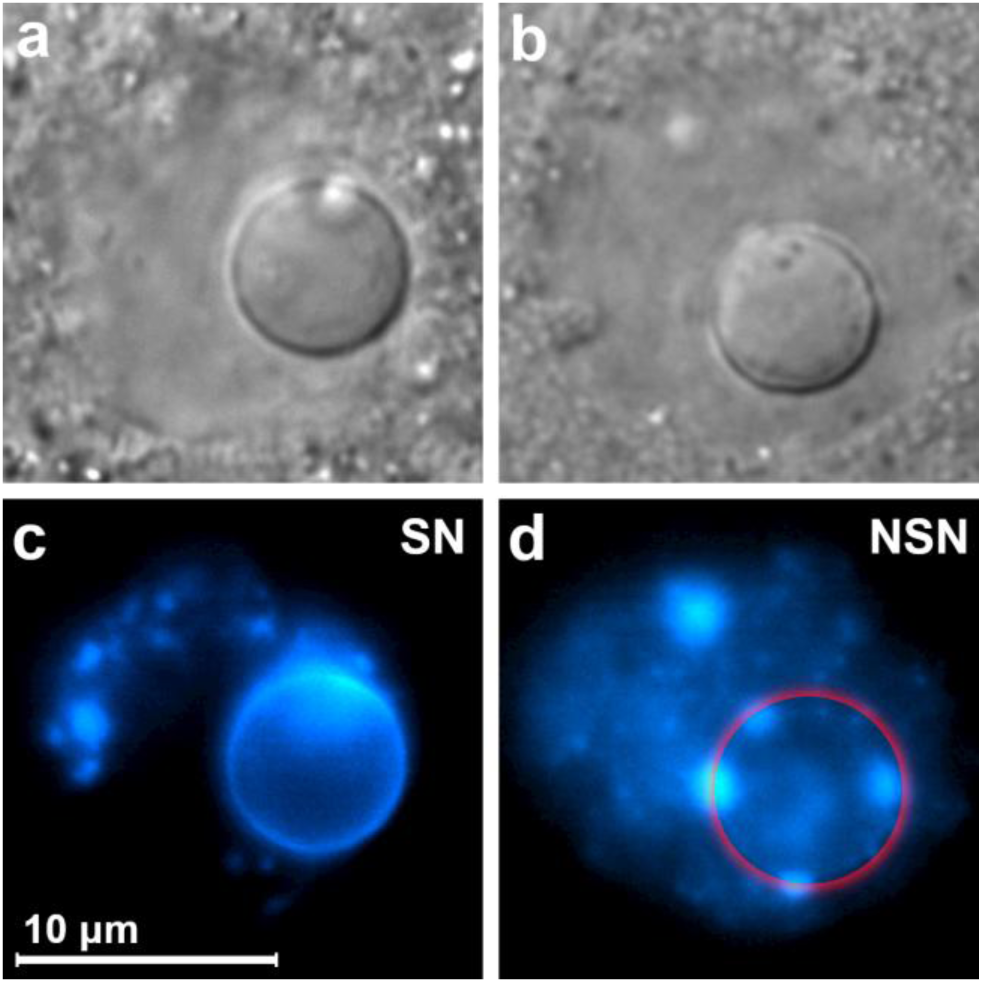
Main oocyte types classified according to chromatin configuration after Hoechst 33342 staining. (**a**, **c**) NSN-type, chromatin is diffusely located in the karyoplasm; (**b**, **d**) SN-type, heterochromatin surrounds the nucleolus with a ring. (**a**, **b**) differential interference contrast microscopy (DIC). (**b**, **d**) Hoechst 33342 staining fluorescent image. Red circle corresponds to nucleolus position.

Subsequent nucleolus displacements to the maximum possible distance in several different directions and its relaxations were observed to reveal the behavior of the entire nucleoplasm volume and classify oocytes according to the types of nucleolus movement and the nature of the chromatin distribution in the nucleus (SN-type or NSN-type). After that, we arrange NSN-oocytes in 4 additional categories in accordance with the specificity of nucleolus motion (Fig. 2).

**Figure 2.**
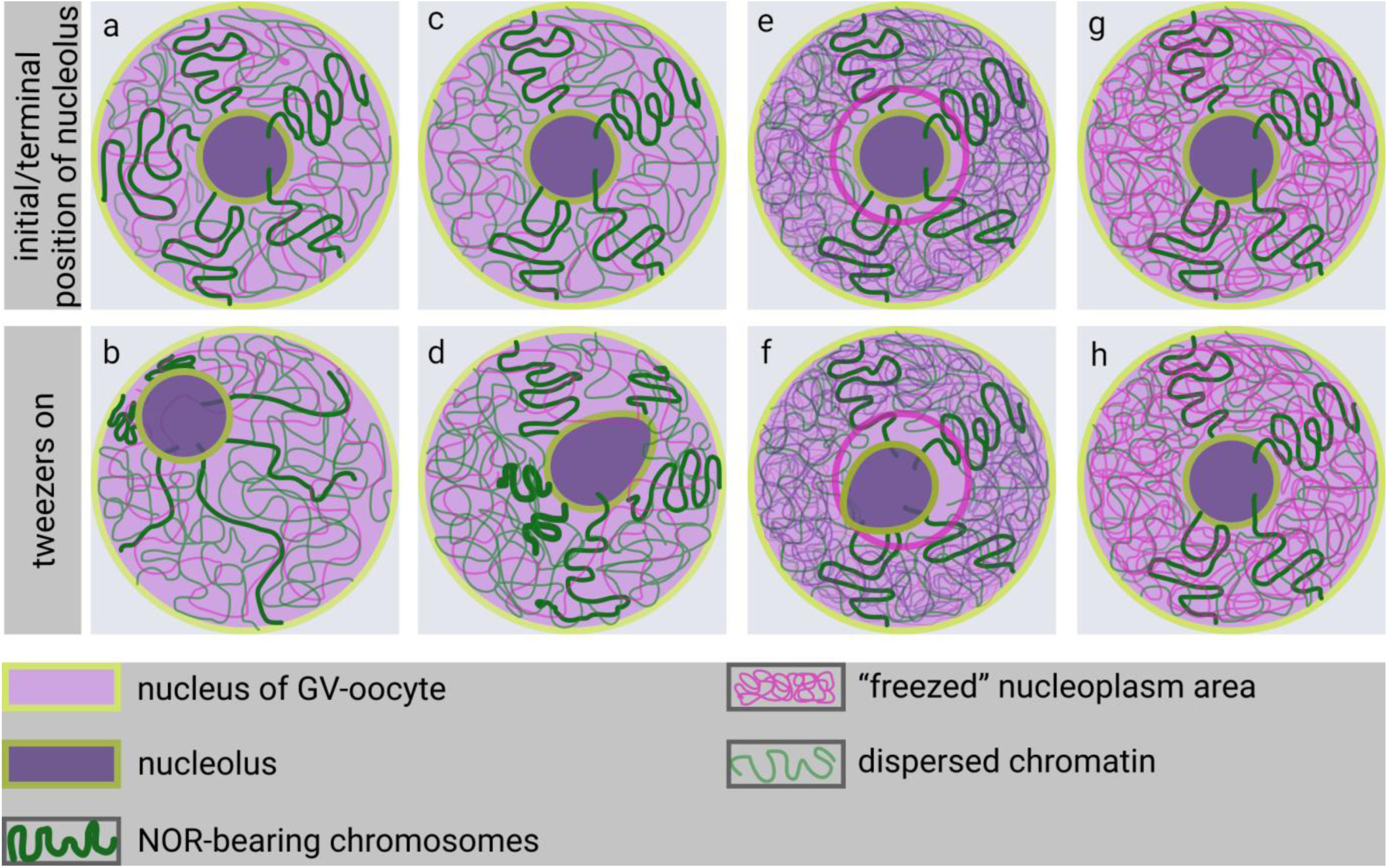
Schematic illustration of 4 types of nucleolar organization and motion within nucleus of NSN-oocytes. **(a**, **b)** the nucleolus freely moves in the space of the nucleus; **(c**, **d)** the nucleolus is slightly displaced and strongly deformed; **(e**, **f)** extremely low mobility, combined with a practically undeformable nucleolus; **(g**, **h)** completely motionless nucleoli.

The phase contrast and fluorescent images of all observed NSN-oocytes were, in fact, the same. Nuclei had the spherical shape, large size and the chromatin was distributed diffusely, but nucleoli moved completely different. Nucleoli of the first and largest group (10 oocytes) freely moved within nucleoplasm in different directions, reached the edge of the nuclear envelope and did not change their shape (Fig. 2a,b).

Nucleoli of the second group (5 oocytes) demonstrated lower and confined motility. Movement of nucleoli was followed by severe deformation into drop-like shape (Fig. 2c,d). Relaxation of nucleoli to initial position was accompanied by relaxation of the nucleoli shape into sphere. Nucleoli reached the edge of nuclear envelope only due to sufficient own deformation. Movements of nucleoli were restricted.

Motion of nucleoli from the third group can be described as extremely limited to very small area, combined with the slight deformation of nucleolar surface by optical trap (Fig. 2e,f). And finally, nucleoli of the fourth group were motionless and embedded into high density nucleoplasmic mesh (Fig. 2g,h). Subsequent nucleolus displacements to the maximum amplitudes revealed biexponential decay response of nucleolus relaxation and following relaxation characteristic times for the first three groups (Table 1).

**Table 1.** Exponential decay times for nucleolus relaxation. Nucleolus was displaced by optical tweezers at a maximum distance from equilibrium position and then relaxed. Tracking data was fitted with biexponential decay function.

| Full mobility<br>(10) | Restricted mobility<br>(5) | Extremely low mobility<br>(6) |
| --- | --- | --- |
| $T_1=0.86\pm0.18$ s | $T_1 = 1.03\pm0.17$ s | $T_1 = 1.13\pm0.15$ s |
| $T_2=6.9\pm0.85$ s | $T_2 = 12.4\pm2.42$ s | $T_2=11.9\pm1.31$ s |

Second mode of displacement was applied to extract the motion of nucleolus without its deformation, possible disruption of large-scale chromatin organization and with the smallest possible amplitude. However high fluctuation rate kept exponential decay fitting away.

## DISCUSSION

Using high-precision optical trapping, we showed that nucleoplasm of mouse oocytes can be investigated by manipulating of the nucleolus as a spherical micron-sized probe. Proposed technique has the potential to extract mesoscale viscosity of nucleoplasm due to the large size (about 5-7 µm) of nucleolus, doesn’t require injection of extra probes, that can shift crowding environment and alter the viscoelastic response of the medium. Series of sequential displacements with several amplitudes of nucleolus revealed anisotropic mechanical properties of NSN-oocytes’ nucleoplasm, despite the seemingly homogenous environment and diffusive chromatin distribution. We described 4 patterns of nucleoplasm-nucleolus interactions, but at this stage didn’t get a clear quantitative differences between these patterns, due to the multiple events and factors involved in nucleolus motion.

Anisotropy of nucleolplasm environment can be attributed to the fact that the oocyte, being a thermodynamically non-equilibrium system, at each point of time undergoes morphogenetic transformations. Thus, there are some topological issues which can be the sources of anisotropy. Up to six NORs are presented in mice genome, involved in nucleolus formation and attached to it.^26^ Concentrated on specific areas of the nucleus, these chromosomes provide asymmetry in the nuclear architecture. Bivalents differ by size, morphology, transcriptional activity, conformation, uneven distribution and position of chiasmata along the homologs. Practically, any act of chromatin remodeling affects motion/position of nuclear body, embedded in it.

Stress-relaxation tests revealed 4 types of interactions between nucleolus and nucleoplasm within NSN-oocyte. Nucleoli of NSN-oocytes from the first group displayed the most dynamic and mobile behavior. NSN-oocytes casually are represented as a non-complete stage of GV-oocyte development. Through the reaching of proper size and global transcriptional silencing, NSN-oocytes, as many authors note, become mature (acquire SN configuration) and after fertilization, develop to the blastocyst stage, but this fact still contradictory.^27-28^ From time-lapse images of large-scale chromatin dynamics of SN-/NSN-oocytes, performed by Belli et al., NSN-to-SN transition is quiet not obvious.^29^ But if we accept the fact, that NSN-configuration is just a step on a way for reaching developmental competence, we can suggest, that relatively high fluidity of nucleoplasm combined with intensive motion of smaller nuclear bodies is supposed to be dominant state in oocytes of this type, corresponding to high rates of transcriptional activity.^30^ From our observations, NSN-oocytes with fluid nucleoplasm is the major population among the whole NSN-oocytes’ pool. Characteristic relaxation times for nucleoli from this group are smaller, than for nucleoli from other groups, especially T2 (2 times smaller). Behavior of nucleoplasm at this state can be described as viscoelastic liquid.

Nucleoli of the second group showed motility constrained on the one side of the GV and combined with nucleolus deformations, but within the same fluid-like nucleoplasm, with intensive motion of smaller nuclear bodies. We suggest that this state is the next phase of transition to SNthrough pSN-configuration (intermediate state) and corresponds emerging chromatin density on the one side of the nucleus (Fig. 3).

**Figure 3.**
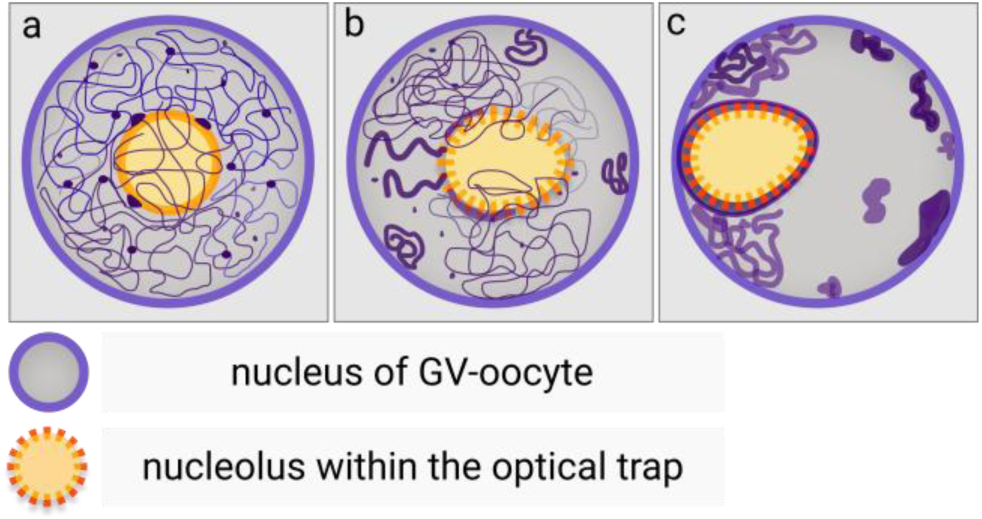
Presumed migration of nucleolus through NSN-to-SN transition. **(a)** nucleolus is centrally located in oocytes of NSN-type; **(b)** due to condensation of NOR-bearing chromosomes, nucleolus migrate to the periphery of nucleus; **(c)** nucleolus is strongly attached to the nuclear membrane via highly condensed chromatin.

Presence of the last two modes remains even more unclear. Nucleoplasm of these NSN-oocytes seemed partly or fully «freezed», despite the fact that chromatin distribution has not changed. Deformation of nucleolus does not occur or expressed weakly, because of highly viscous and stiff environment. Own nucleolar diffusive motion was also suppressed. This nucleoplasm state can be described as viscoelastic solid. Initially, we assume that this state was caused by triggering of cell culture/experimental conditions (e.g. osmotic stress or temperature fluctuations) or oocyte atresia. But, as was shown, osmotic challenge,^31^ temperature fluctuations stimulate changes of chromatin condensation, which wasn’t observed after Hoechst staining and fluorescent imaging – distribution of chromatin corresponded to NSN-type configuration and cytoplasm of several oocytes demonstrated intensive motion of cytoplasmic bodies/vesicles. Some studies reported that «freezing» occurs in cells with pathological processes and characterized by gelation of cellular interior due to formation of protein/RNA aggregates.^32^ Thus, by using optical trapping, we were able to observe various states of possible nuclear matrix (presence of nuclear matrix is still debated)^33-34^ within oocytes of the same meiotic phase, chromatin distribution, under the equal culturing conditions *in vivo*.

Averaged relaxation characteristic times turned out to be particularly the same for 3 patterns of nucleolar movement, despite the fact that images, obtained during the experiment, indicated significant variety. To estimate this differences correctly, relaxation times should be compared specifically between similar directions or chromosomes. Differences between patterns should be considered blurry; in some cases, we met mixed types of chromatin configuration. nucleolar movement.

Biexponential decay relaxation was extracted for both types of stress-relaxation regimes. In some cases, relaxation started as exponential but soon after affected by fluctuations with considerable amplitudes (Fig. 4). Relaxation curves for the small amplitudes in general were also characterized by biexponential decay as well, but haven’t been fitted correctly due to high fluctuation rate.

**Figure 4.**
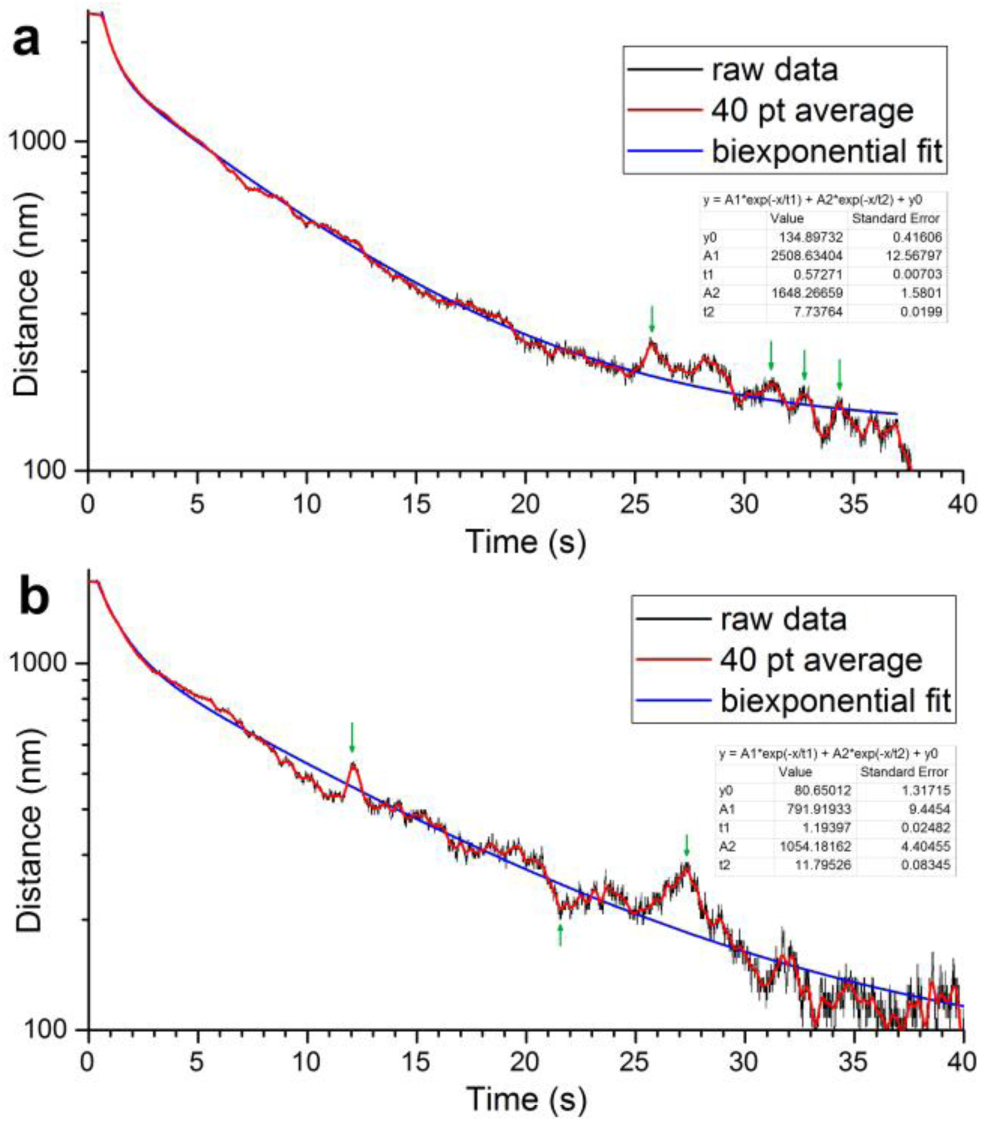
Curves represent relaxation of nucleoli after displacement to the maximal amplitude in the second group of NSN-oocytes with restricted mobility. Green arrows correspond to fluctuations emerged throughout relaxation.

Previously, it was shown that motion of nuclear bodies (such as Cajal bodies or PML bodies) is confined by the presence of corrals, the boundaries formed by adjacent chromatin masses ^35^. Configuration and viscosity to elasticity ratio of adjacent chromatin determine type of motion experienced by nuclear bodies. Initiating very small displacement of nucleolus, we, potentially, just «deform» corral. When the optical trap was turned off, the nucleolus relaxed and after momentary contact with the opposite surface of the corral, bounced off it and relaxed again. This situation illustrates elastic component of nucleus environment. In the regime, when nucleolus displacement reached its maximum, corral structure was destroyed by the forces, generated by optical trapping and nucleolus was taken off the corral and nucleoplasm material experienced plastic deformation. Nucleolus relaxed to its initial point, pushing through chromatin masses, but sometimes did not return to the starting position, especially after several displacements, probably, due to remodeling of chromatin, attached to it. The size of nucleolus is quiet large and the only obstacle of comparable size that could significantly affect its movement is chromatin and its spatial distribution, because other nuclear bodies are much smaller.

Definitely, kinetics of nucleoli relaxation cannot be explained simply by topological or spatial changes of nucleoplasm material. Chromatin and nuclear bodies of NSN-oocytes during diplotene stage are exposed to forces, generated by key molecular motors, such as topoisomerases, RNA-polymerases, gyrases. Nucleoli of the NSN-type nuclei are also transcriptionally active; rRNA-polymerases, involved in transcription, affect own nucleolar motion. Studies of Cajal bodies and PML bodies motility revealed that nuclear bodies experience 4 types of motion depending on adjacent chromatin organization.^35^ Forces, created by cytoskeleton and motor proteins during oocyte growth also contribute into chromatin dynamics through telomeres, attached by SUN/KASH complex on nuclear envelope. Well known, that active motion of telomeres or subtelomeric regions promotes pairing of homologs during early prophase,^36^ but in pachytene telomeres motion is inhibited in mice.^37^ It can be assumed, that for further segregation of chromosomes around the nucleolus (SN-stage) during diplotene/diakinesis, revival of telomeres movement is essential.

With further research, differences in nucleoplasmic mobility may be a key for understanding the reasons, why NSN-oocytes delay meiosis compared to SN-oocytes, or identifying a population of oocytes, that, naturally, cannot complete meiosis due to an altered micromechanics of nucleoplasm. Nucleoli of maturing oocytes are considered as a points of chromosomes aggregation to provide a «precursor» of metaphase plate and resolve meiotic maturation.^26^ Probably, in oocytes with dense nucleoplasm, moving of single chromosomes to nucleolus can be inhibited by highly viscous/dense environment and GVBD is retarded as well or does not occur.

## CONCLUSION

We have illustrated a technique of the nucleolus manipulation in the nucleoplasm performed by optical trapping. Using the nucleolus as a microprobe makes this technique minimally invasive. The relaxation times of the nucleolus under the force of optical tweezers showed anisotropic properties of the nuclear material. Relatively large size of nucleolus facilitated extracting properties of whole nuclear volume and revealed 4 different patterns of nucleolar motion, providing unique model system. These observations cannot be reached by employment of smaller nuclear bodies or by injection of particles due to higher risks of cellular interior disruption. However, unraveling more specific causes of anisotropy and nucleoli motion requires additional research, since oogenesis is a complex, multifactorial process and aspects of the nucleolus motility are restricted by its own biophysical properties as well as the properties of its highly crowded microenvironment.

## MATERIALS & METHODS

### Collection of oocytes

Female mice (C57BL/CBA) aged 6-9 weeks were injected with 7.5 IU PMSG (pregnant mare’s serum gonadotropin). After 48 hours, oocytes were collected from ovaries by puncturing follicles with a needle in M2 medium and then cleaned from cumulus cells with hyaluronidase. To perform stress-relaxation tests every single oocyte was transferred into individual 30 μl drop of M2 medium on the coverslip. The work was carried out on a temperature controlled microscope stage. A total of 65 oocytes were examined. Of these, 28 oocytes had NSN-configuration of chromatin.

### Laser setup

Optical trapping of nucleoli was performed by continuous wave radiation from a Ti:Sapphire laser (wavelength 790 nm). Laser radiation was coupled to a microscope (Olympus IX71) and focused by an objective lens (60x 0.7 NA). Laser power at the focus of the objective lens was kept at 280 mW, which corresponds to 10 pN maximal transverse gradient force, applied to a nucleolus in the optical trap and measured by calibrated flow in microfluidics system with oocyte. Stokes’ drag force method was used to calibrate optical tweezers. Oocyte was injected into transparent microchannel with the depth of 200 μm. The force applied on a trapped oocyte was balanced with the viscous drag force, generated by phosphate-buffered saline constant flow.

### Imaging

Series of stress-relaxation tests on nucleolus were visualized by differential interference contrast microscopy (DIC) and were recorded by high-speed camera at 100 fps (Ximea xiC MC023MG-SY). To visualize chromatin structure, oocytes were treated with 1 mg/ml Hoechst 33342 in M2 medium for 10 minutes. Chromatin visualization was performed after the series of nucleolus displacements to avoid changing of nucleoplasm viscoelastic response.

### Nucleolus tracking

The recorded videos were processed using custom-made software, which provided detection of the nucleolus center two-dimensional coordinates with subdiffractional resolution. Mathematical algorithm of coordinates detection was performed through scanning of template across the image with subsequent matching of template with the image for each frame. Resolution was mathematically increased and it was equal 1/16 of the image pixel size or approximately 3 nm.

### Optical trapping

When the laser was turned on, the nucleolus was pulled into the optical trap and could be moved within the nucleoplasm space. Stress-relaxation tests of the nucleolus were performed sequentially in different directions with an interval of several tens of seconds. Immediately after the relaxation of whole nucleus, chromatin structure was observed (stained with Hoechst 33342).

### Mathematical analysis

Relaxation curves exponential fitting were performed by OriginPro software (Origin Lab Corporation).

## ABBREVIATIONS

GV: germinal vesicle,
FCS: fluorescence correlation spectroscopy,
FRAP: fluorescence recovery after photobleaching,
NSN: non-surrounded nucleolus,
SN: surrounded nucleolus,
GVBD: germinal vesicle breakdown,
NOR: nucleolus organizer region.

## ACKNOWLEDGEMENTS

This work was supported by the Russian Foundation for Basic Research (RFBR) Grant No. 18-33-01080. Experiments were performed using the equipment of Semenov FRCCP RAS CCE (no. 506694).

